# Investigating the Role of Hypoxia-Induced Migration in Glioblastoma Growth Rates

**DOI:** 10.1101/862920

**Authors:** Lee Curtin, Andrea Hawkins-Daarud, Kristoffer G. van der Zee, Kristin R. Swanson, Markus R. Owen

## Abstract

We analyze the wave-speed of the Proliferation Invasion Hypoxia Necrosis Angiogenesis (PIHNA) model that was previously created and applied to simulate the growth and spread of glioblastoma (GBM), a particularly aggressive primary brain tumor. We extend the PIHNA model by allowing for different hypoxic and normoxic cell migration rates and study the impact of these differences on the wave-speed dynamics. Through this analysis, we find key variables that drive the outward growth of the simulated GBM. We find a minimum tumor wave-speed for the model; this depends on the migration and proliferation rates of the normoxic cells and is achieved under certain conditions on the migration rates of the normoxic and hypoxic cells. If the hypoxic cell migration rate is greater than the normoxic cell migration rate above a threshold, the wave-speed increases above the predicted minimum. This increase in wave-speed is explored through an eigenvalue and eigenvector analysis of the linearized PIHNA model, which yields an expression for this threshold. The PIHNA model suggests that an inherently faster-diffusing hypoxic cell population can drive the outward growth of a GBM as a whole, and that this effect is more prominent for faster proliferating tumors that recover relatively slowly from a hypoxic phenotype.

## 1 Introduction

Glioblastoma multiforme (GBM) is the highest grade of glioma from the World Health Organization and is the most aggressive type of primary brain tumor [9]. It is uniformly fatal with an average survival time from diagnosis of only 15 months with standard of care treatment [14]. The standard therapy regime for this disease is a combination of resection, radiation and chemotherapy [14, 15]. Magnetic Resonance Imaging (MRI) is the standard imaging modality for GBMs and is used routinely to monitor tumor growth and development throughout the progression of the disease. Different MRI sequences such as gadolinium-enhanced T1-weighted (T1Gd) and T2-weighted (T2) are used to identify the gross tumor volume. T1Gd shows gadolinium that has leaked into brain tissue, and T2 shows water that has done the same, which is known as edema. However, these MRI sequences together do not show a complete picture. Infiltrating tumor cells also exist beyond the resolution of these MRI sequences. In fact, malignant glioma cells have been cultured from histologically normal healthy tissue at a distance of 4cm from the gross tumor volume identified by MRI scans [13].

Hypoxia has been shown to induce more migration in glioma cells [7, 23]. There is also evidence that glioma cells follow a dichotomy of migration and proliferation [3] and evidence of a lower proliferation marker for cells that exist in hypoxic regions of GBMs [1]. Tumors in hypoxic conditions release angiogenesis-promoting factors to encourage vessels to grow towards them and provide nutrients [4, 8, 24]. This process also occurs in normoxic conditions at a lower level [24]. Necrosis occurs in the vast majority of GBMs and presents in the core of the tumor [9].

Over the past 20 years, there have been many partial differential equation models that simulate GBM cell density and have provided various insights into this disease [6, 10, 11, 16–19, 19–21]. One such model is the Proliferation Invasion Hypoxia Necrosis Angiogenesis (PIHNA) model, which has been used to analyze the mechanistic properties of GBMs that lead to observed imaging features and has shown similar growth and progression patterns to those seen in patient tumors [20]. We carry out a traveling wave analysis on the PIHNA model to determine which parameters drive the outward growth of the tumor as a whole, and compare these analytical predictions with computational simulations in the cases of varying relative rates of migration between hypoxic and normoxic tumor cells. We find that the normoxic cell migration and proliferation rates, *D*_*c*_ and *ρ*, respectively, drive the minimum wave-speed in the PIHNA model, which is given by

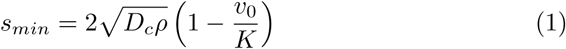

and also depends on the initial background vasculature in the model, *v*_0_, relative to the spatial carrying capacity, *K*. We find that *s*_*min*_ holds for published results using the PIHNA model as they have not allowed for different hypoxic cell and normoxic cell migration rates. We allow these migration rates to be different in the model and observe the effect of this variability on simulated tumor growth rates. We find that a faster-than-minimum wave-speed is achieved when hypoxic cells migrate sufficiently faster than normoxic cells and find a threshold above which these dynamics can occur. This threshold depends on the proliferation rate *ρ*, the switching rate back from hypoxic cells to normoxic cells *γ*, and *v*_0_*/K*. We denote this threshold *k*, and it is given by

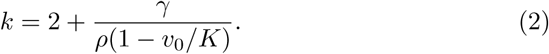

These results are then confirmed and explored computationally through further model simulations.

We introduce the PIHNA model in the next section before calculating the expression for the minimum wave-speed in Section 3. Following this, in Section 4 we find the threshold, *k*, on the relative migration between hypoxic and normoxic cells under which the minimum wave-speed is achieved. We then move onto PIHNA simulations in Section 5 to computationally validate our findings.

## 2 The PIHNA Model

The PIHNA model [20] simulates five different species and their interactions:

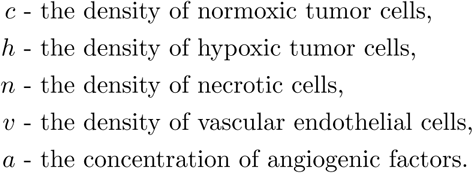

The dimensions of *c, h, v* and *n* are cells/mm^3^ of tissue. The angiogenic factor, *a*, is a diffusing concentration with dimensions *µ*mol/mm^3^ tissue.

Normoxic cells proliferate with rate *ρ* and migrate with rate *D*_*c*_, whereas hypoxic cells do not proliferate but migrate with rate *D*_*h*_. Cells convert between normoxic and hypoxic phenotypes depending on the ability of the local vascular density to provide nutrients at their location; hypoxic cells in the model become necrotic if they remain in such a region. When any other cell type meets a necrotic cell, they become necrotic with rate *α*_*n*_. Previous publications on the PIHNA model have set the migration rate of hypoxic cells to be equal to that of normoxic cells, such that *D*_*h*_ = *D*_*c*_. However, hypoxia has been shown to promote GBM cell migration, so we have allowed for this to be varied in the PIHNA model [7, 23].

Angiogenic factors are created by the presence of normoxic and hypoxic cells, decay naturally and are consumed through the creation and presence of vascular cells (*v*). Angiogenic factors are only consumed by vasculature and not tumor or necrotic cells. Necrotic cells are dead cells and their degradation is not considered in the model.

The governing partial differential equations for the PIHNA model are

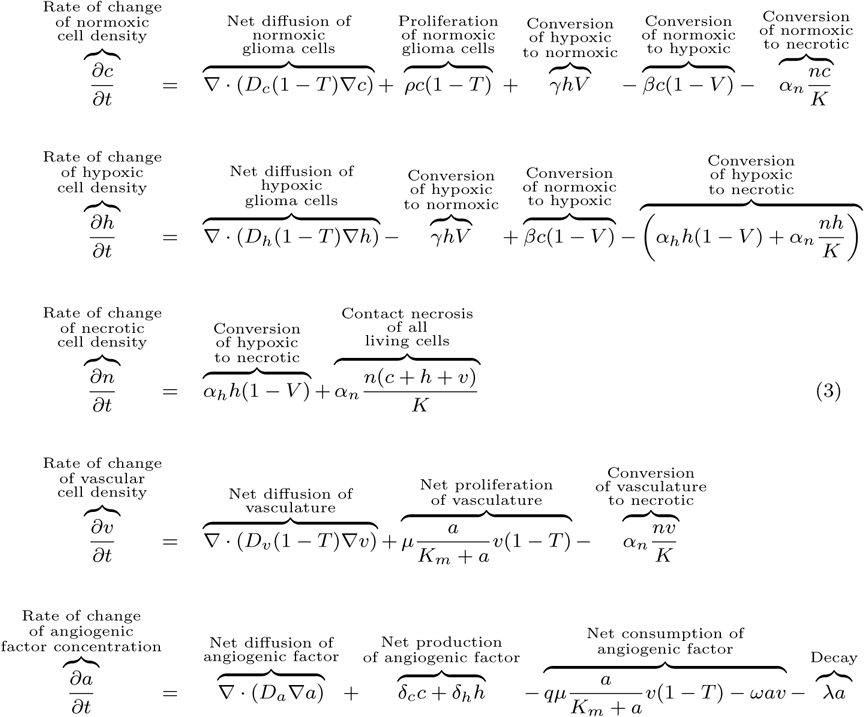

where

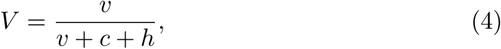

and

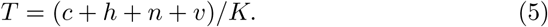

The term *V* models the relationship between the vasculature and its effect on the tumor. Note that *V* take values in [0, 1] such that it affects the switching rates between the populations *c, h* and *n*. A value of *V* (*c, h, v*) ≈ 0 corresponds to a very inefficient vasculature that cannot provide sufficient nutrients to the local tumor population; this would increase the conversion of normoxic cells to hypoxic cells and in turn necrotic cells. A high *V* (*c, h, v*) ≈ 1 promotes a normoxic phenotype. It is worth noting that, once necrotic cells are present in a simulation, they will always increase in population due to the contact necrosis in the model.

The expression for *T* defined in Equation (5) is a spatiotemporal measure of the relative density of the cells in a region. It is used to limit growth and migration and used as a threshold to determine which densities would appear on different MRI sequences. Substituting Equation (3) into Equation (5) gives

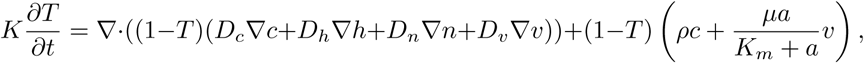

from which it is clear that at *T* = 1 the reaction and diffusion terms vanish, which implies *T* is restricted by the upper bound of 1 (as long as *T* (*x*, 0) ≤ 1). As *T* is a sum of non-negative components and *K* > 0, we have that *T* ≥ 0. Therefore, we have that *T* ∈ [0, 1] for sufficient initial conditions, for all *x* and *t* ≥ 0.

Following the literature, we have assumed that a total relative cell density of at least 80% is visible on a T1Gd MRI, and a total relative density of at least 16% is visible on a T2 MRI [17, 20]. In the PIHNA model, this translates to *T* ≥ 0.8 being visible on T1Gd MRI and *T* ≥ 0.16 being visible on T2 MRI. By construction the T1Gd radius is always less than or equal to the T2 radius, which agrees with patient data [5].

For the purposes of the wave-speed calculations, we consider the PIHNA model in a one-dimensional spherically symmetric case with zero-flux boundary conditions at the end points, *r* = 0 and *r* = *r*_*end*_. This does not take into account the full anatomy of the brain, but it is useful to gain insight into the behavior of the PIHNA model. The initial condition is given by

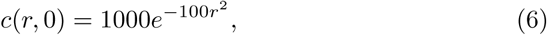

to simulate a small initiating population of normoxic tumor cells decreasing away from the core of the tumor. We also have *h*(*r*, 0) = 0, *n*(*r*, 0) = 0, *v*(*r*, 0) = 0.03*K* and *a*(*r*, 0) = 0. We run the PIHNA simulations with the parameters found in Table 1.

**Table 1.** Parameter definitions and values for the PIHNA model. A justification of parameters can be found in the supplementary material of [20]. *We have altered these rates in this formulation of PIHNA, which have not been changed previously.

|  | Definition | Value/Range | Units | Source |
| --- | --- | --- | --- | --- |
| $D_c$ | Diffusion rate of normoxic cells | 10 – 100 | $\frac{\text{mm}^2}{\text{year}}$ | [5] |
| $D_h$ | Diffusion rate of hypoxic cells | $(0.1 - 100)D_c$ | $\frac{\text{mm}^2}{\text{year}}$ | [5, 10]* |
| $\rho$ | Proliferation rate of normoxic cells | 10 – 100 | 1/year | [5] |
| $\beta$ | Switching rate from normoxia to hypoxia | $0.1\rho$ | 1/year | [20] |
| $\gamma$ | Switching rate from hypoxia to normoxia | 0.005 – 0.5 | 1/day | [20]* |
| $\alpha_h$ | Switching rate from hypoxia to necrosis | $0.5\beta$ | 1/year | [20] |
| $\alpha_n$ | Rate of contact necrosis | $\log(2)/50$ | 1/day | [12] |
| $D_v$ | Diffusion rate of endothelial cells | 0.18 | $\frac{\text{mm}^2}{\text{year}}$ | [20] |
| $D_a$ | Diffusion rate of angiogenic factors | 3.15 | $\frac{\text{mm}^2}{\text{year}}$ | [20] |
| $\delta_c$ | Normoxic cell production rate of angiogenic factors | $2.77 \times 10^{-13}$ | $\frac{\mu\text{mol}}{\text{cell} \times \text{year}}$ | [20] |
| $\delta_h$ | Hypoxic cell production rate of angiogenic factors | $5.22 \times 10^{-10}$ | $\frac{\mu\text{mol}}{\text{cell} \times \text{year}}$ | [20] |
| $\mu$ | Angiogenesis vasculature production rate | $\log(2)/15$ | 1/day | [20] |
| $q$ | Consumption of angiogenic factors per cell | 1.66 | $\mu\text{mol}/\text{cell}$ | [20] |
| $\lambda$ | Natural decay rate of angiogenic factors | 15.6 | 1/day | [20] |
| $\omega$ | Rate of removal of angiogenic factors by vasculature | $\lambda/v_0$ | $\frac{1}{\text{cell} \times \text{year}}$ | [20] |
| $K$ | Maximal cell density | $2.39 \times 10^8$ | cells/cm<sup>3</sup> | [20] |

In all simulations, the tumor and necrotic cell densities spread outwards. A peak in normoxic cell density leads and is followed by a peak in hypoxic cell density and then a zone of necrosis, as can be seen in Figure 1; this figure also shows how we calculate the wave-speed values from simulations.

**Fig. 1.**
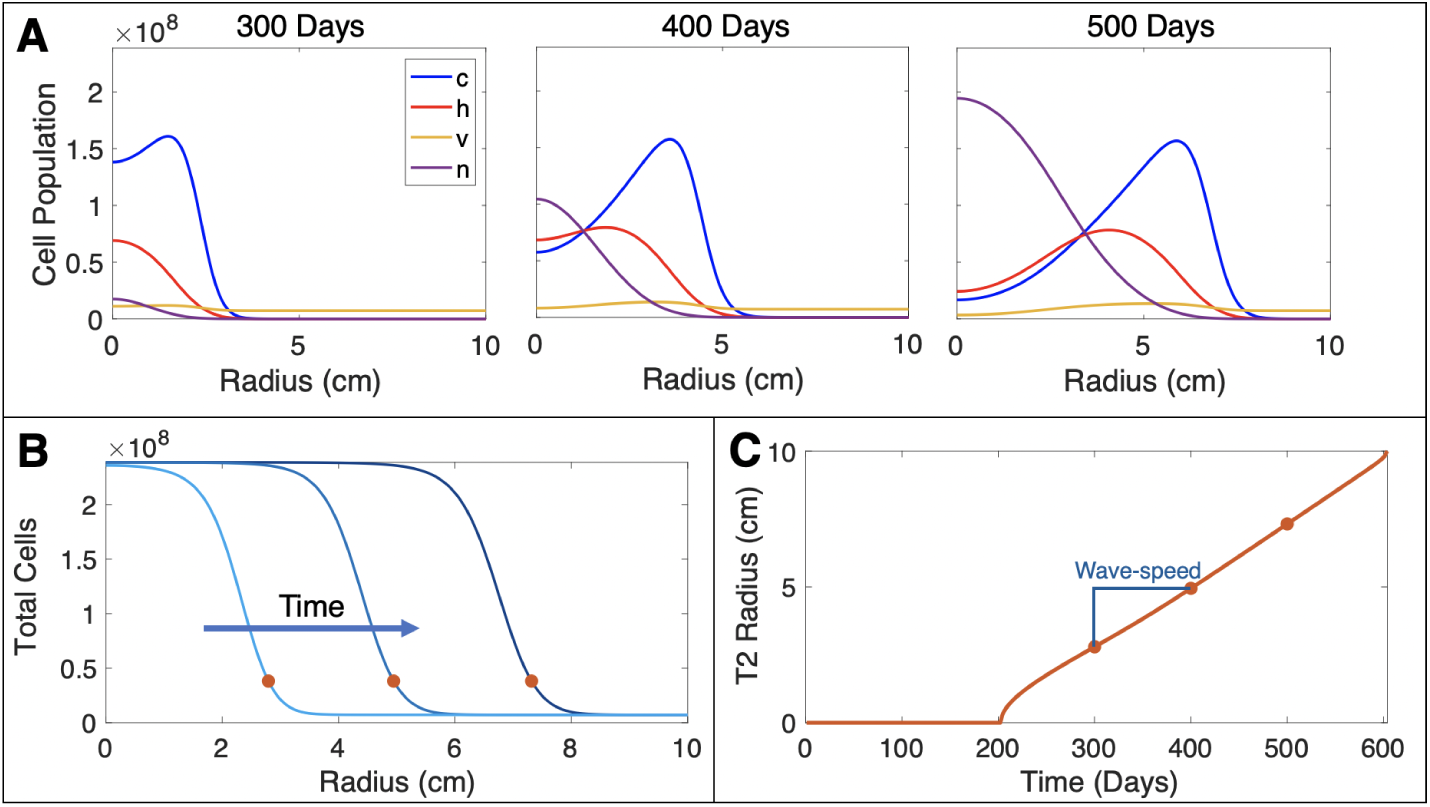
(A) Three example time points (300 days, 400 days and 500 days) of a simulation with *D*_*c*_ = 10^1.5^mm^2^/year, *ρ* = 10^1.5^/year, *γ* = 0.05/day and *D*_*h*_*/D*_*c*_ = 10. All cell types are shown and move outwards over time. Necrosis develops in the core of the tumor. (B) Total cells over time for the three time points shown in Subfigure A. The dots correspond to the T2 radius at each time point. (C) The T2 radius shown over time for the same simulation. This radial growth is non-linear for small tumor sizes, but settles to a linear rate, which is the wave-speed of the simulation.

## 3 Minimum Wave-speed for the PIHNA Model

In a similar fashion to the well-established minimum wave-speed of Fisher’s Equation [2] that has been used for the Proliferation Invasion tumor growth model [17], we carried out a wave-speed analysis to find an analytical expression for the tumor wave-speed in the PIHNA model. Note that in spherically symmetric coordinates, the wave-speed asymptotically approaches that of a planar wave-speed. We start by linearizing the model ahead of the leading edge of the wave, that has the initial condition of (*c, h, n, v, a*) = (0, 0, 0, *v*_0_, 0); this gives an expression of

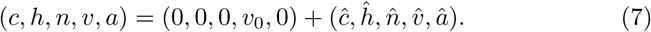

Substituting Equation (7) into the PIHNA model (Equation (3)) and discarding non-linear terms leads to the following set of equations:

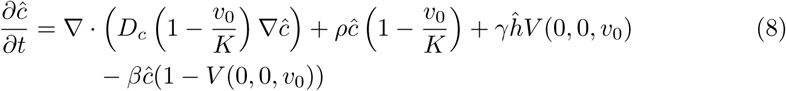

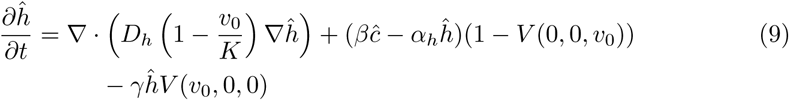

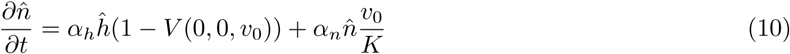

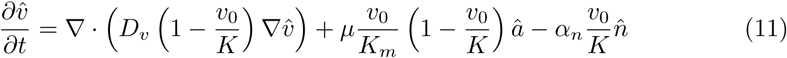

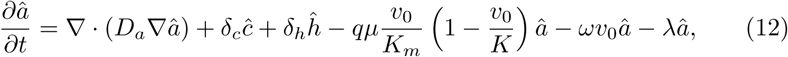

The equations for *ĉ* and *ĥ* decouple from the system. We will analyze these two equations to look for traveling wave solutions of the form

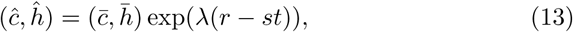

where *s* is the wave-speed. Substituting Equation (13) into Equations (8) - (9), gives rise to the following equations

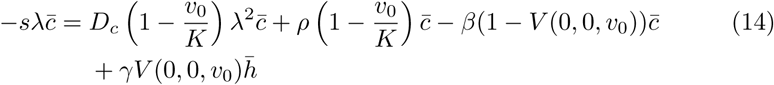

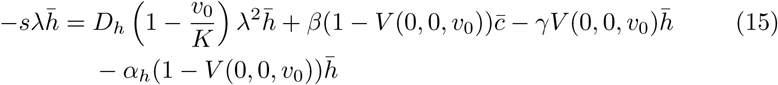

We rearrange the equations into a matrix form given by

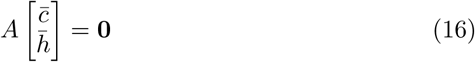

and note the determinant of *A*, det(*A*), needs to be zero in order to give non-trivial solutions. Setting det(*A*) = 0 leads to

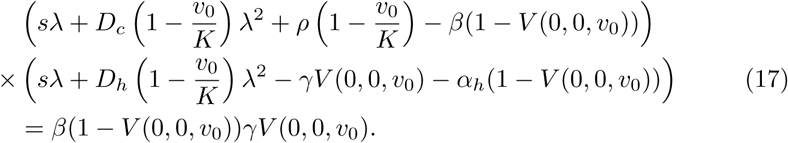

We note that the vasculature ahead of the wave is as efficient as possible in the absence of tumor cells, giving *V* (0, 0, *v*_0_) = 1; this leads to cancellation in Equation (17), which becomes

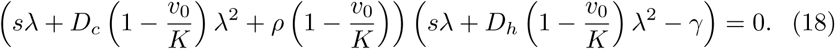

We can then find the eigenvalues for Equation (18) as functions of the wave-speed, *s*. We have four eigenvalues, given by

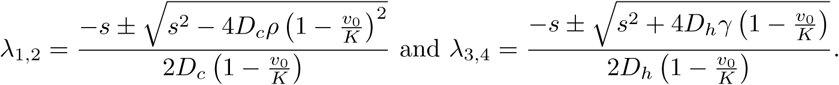

We have also found the corresponding eigenvectors for all of our eigenvalues, which we shall denote *V*_*i*_ for each *λ*_*i*_. These are given by the following expressions, up to a proportional constant:

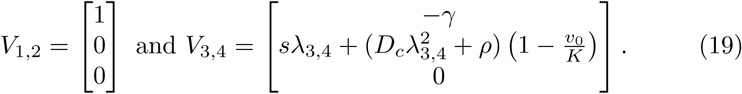

The terms *λ*_1, 2_ are both negative as *s* > 0 by assumption. Due to positive restrictions on the state space (negative populations do not make any biological sense), a spiral approach around the point (0, 0, 0, *v*_0_, 0) cannot occur. Therefore, we need the discriminant of the set of quadratic *λ*_1, 2_ solutions to be greater than or equal to zero. In other words,

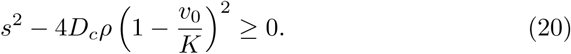

Therefore, we have a minimum wave-speed of

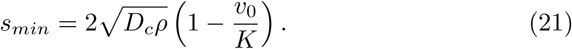

There is no minimum wave-speed associated with the eigenvalues *λ*_3,4_. However, *λ*_4_ can play a role in the wave-speed of the model; we explore this in the next section.

## 4 Conditions for Faster Wavespeeds

In the previous section, we found an expression for the minimum wave-speed in the PIHNA model. The PIHNA model will follow this minimum wave-speed if the eigenvalue *λ*_1_ evaluated at this minimum gives the smallest possible negative eigenvalue of *λ*_1,2,3,4_. If there exists some *s* > *s*_*min*_ such that 0 > *λ*_*i*_(*s*) > *λ*_1_(*s*_*min*_) for some *i* = 1, 2, 3, 4, we will see the emergence of a solution with a larger wave-speed. In this section we will compute a threshold below which the minimum wave-speed is achieved but above which there can be other dynamics emerging.

We will call each eigenvalue evaluated at 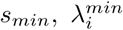, for *i* = 1, *…*, 4. We start by noting that *λ*_2_ ≤ *λ*_1_ and *λ*_3_ > 0, so neither of those can be negative with a smaller magnitude than 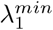 to change the PIHNA wave-speed dynamics. As *λ*_4_ becomes less negative for increasing values of *D*_*h*_, there is a threshold value of *D*_*h*_*/D*_*c*_ that leads to 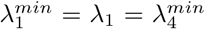 for which the minimum wave-speed is still achieved. For values of *D*_*h*_*/D*_*c*_ that are smaller than this threshold, the minimum wave-speed will still be achieved. However, larger values of *D*_*h*_*/D*_*c*_ may lead to a faster wave-speed, as the eigenvalues become smaller than 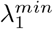. We have

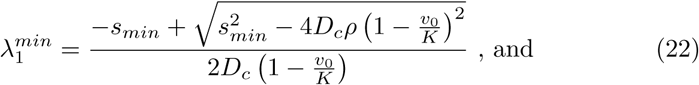

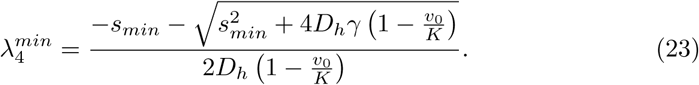

Setting 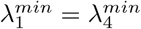 and using our expression for *s*_*min*_ (Equation (21)) leads to

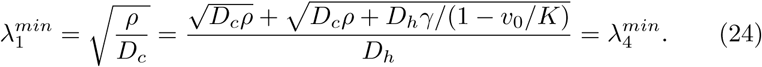

Solving for *D*_*h*_*/D*_*c*_ gives the non-trivial solution

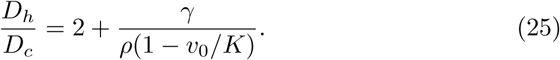

We will define this threshold of *D*_*h*_*/D*_*c*_ as *k*, so we have

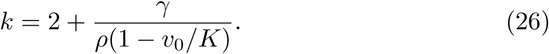

Note that as *v*_0_ ≤ *K, k* ≥ 2. So for the faster wave-speeds to occur, we need the hypoxic cell migration rate to be at least twice as fast as the normoxic cell migration rate. For *D*_*h*_*/D*_*c*_ = 1, as is the case in previous PIHNA publications, we do not expect faster wave-speeds to occur, regardless of other simulation parameters.

## 5 Simulation Results

To calculate the simulated wave-speed in numerical simulations, we thresholded the total cell density at *T* = 0.16, which is a commonly assumed cell density threshold for visible tumor on T2 MRI [17]. Following the establishment of a wave front, the simulated wave-speed levels out to a fixed value, see Figure 1. We analyze the wave-speed of large tumors to ensure we are analyzing established wavefronts while minimizing numerical error. We are particularly interested in the effect on the wave-speed of varying hypoxic cell migration rates, more specifically the change in their migration rate compared with normoxic cells (*D*_*h*_*/D*_*c*_), which has been allowed to vary in the PIHNA model for the first time. Numerical simulations are run on a spherically symmetric domain, with a step size of 0.01mm. All simulations were run in Matlab 2018a using the inbuilt solver *pdepe.m*.

### 5.1 Relatively Fast Hypoxic Cell Diffusion Rates Increase Wave-speed

The wave-speed for PIHNA simulations with *D*_*h*_*/D*_*c*_ ≤ *k* converges towards *s*_*min*_. However, if we compute the wave-speed for simulations where *D*_*h*_ > *kD*_*c*_, we see that the wave-speed can be faster, and continues to increase for larger *D*_*h*_*/D*_*c*_ values; an example of this can be seen in Figure 2. Computing the corresponding eigenvalues shows a change in behavior for values of *D*_*h*_*/D*_*c*_ > *k*. We also plot *k* on Figure 2, in which case *k* = 2.60 (three significant figures). These values of *D*_*c*_ and *ρ* are biologically realistic and based on the mean of previous migration and proliferation rate estimates from a similar mathematical model of GBM growth and patient-specific MRI data [22].

**Fig. 2.**
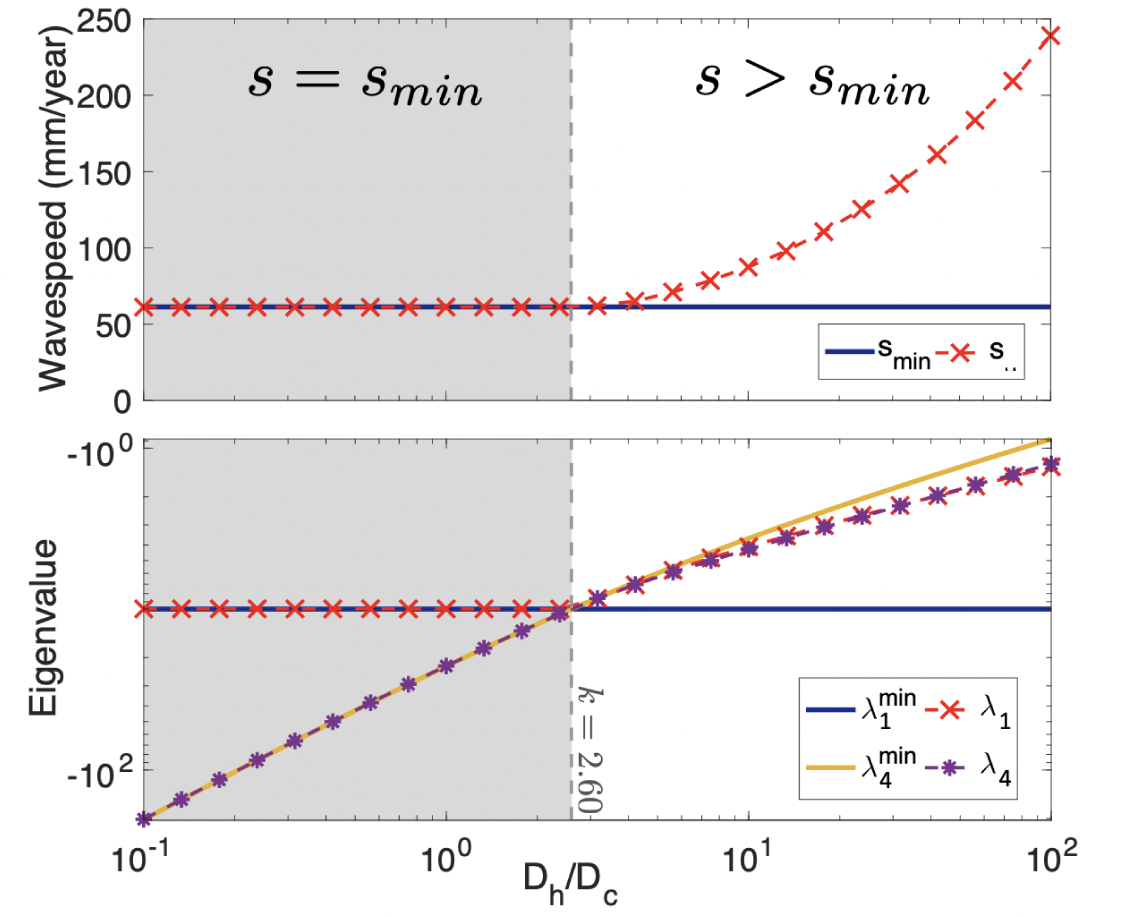
As *D*_*h*_*/D*_*c*_ is increased with *D*_*c*_ = 10^1.5^mm^2^/year, *ρ* = 10^1.5^/year and *γ* = 0.05/day, we see an increase in the converged numerical wave-speed past the threshold on *D*_*h*_*/D*_*c*_ of *k* = 2.60. Wave-speeds taken as the average speed between 8 and 8.5cm of growth on simulated T2 MRI (16% total cell density threshold).

From these observations and our analysis in Section 4, we can deduce that if the hypoxic cell migration is sufficiently faster than the normoxic cell migration (such that *D*_*h*_*/D*_*c*_ > *k*), the hypoxic cell population drives the outward growth of the tumor in the PIHNA model. This behavior intuitively agrees with the biological cell movement patterns that the model is trying to capture; cells moving faster dominate the growth outwards as they search for nutrients.

Focusing on the eigenvectors corresponding to the least negative eigenvalues, *V*_1_ and *V*_4_, we see that they influence the dynamics of the model. By plotting the normoxic cell density across space against the hypoxic cell density across space for a fixed time point where each simulation has converged to a stable wave-speed, together with *V*_1_ and *V*_4_, we can see how the traveling wave trajectory approaches the state ahead of the wave front. We present two simulations with their corresponding *V*_1_ and *V*_4_ eigenvectors in Figure 3, one for *D*_*h*_*/D*_*c*_ = 10^−1^ and another for *D*_*h*_*/D*_*c*_ = 10^1^. For *D*_*h*_*/D*_*c*_ = 10^−1^, where the wave-speed follows the predicted minimum value, we see that the model approaches along (*c, h*) = (0, 0) along the eigenvector *V*_1_, whereas for *D*_*h*_*/D*_*c*_ = 10^1^, the approach is along *V*_4_. In the linearized regime, we expect that 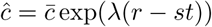, such that

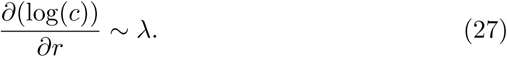

**Fig. 3.**
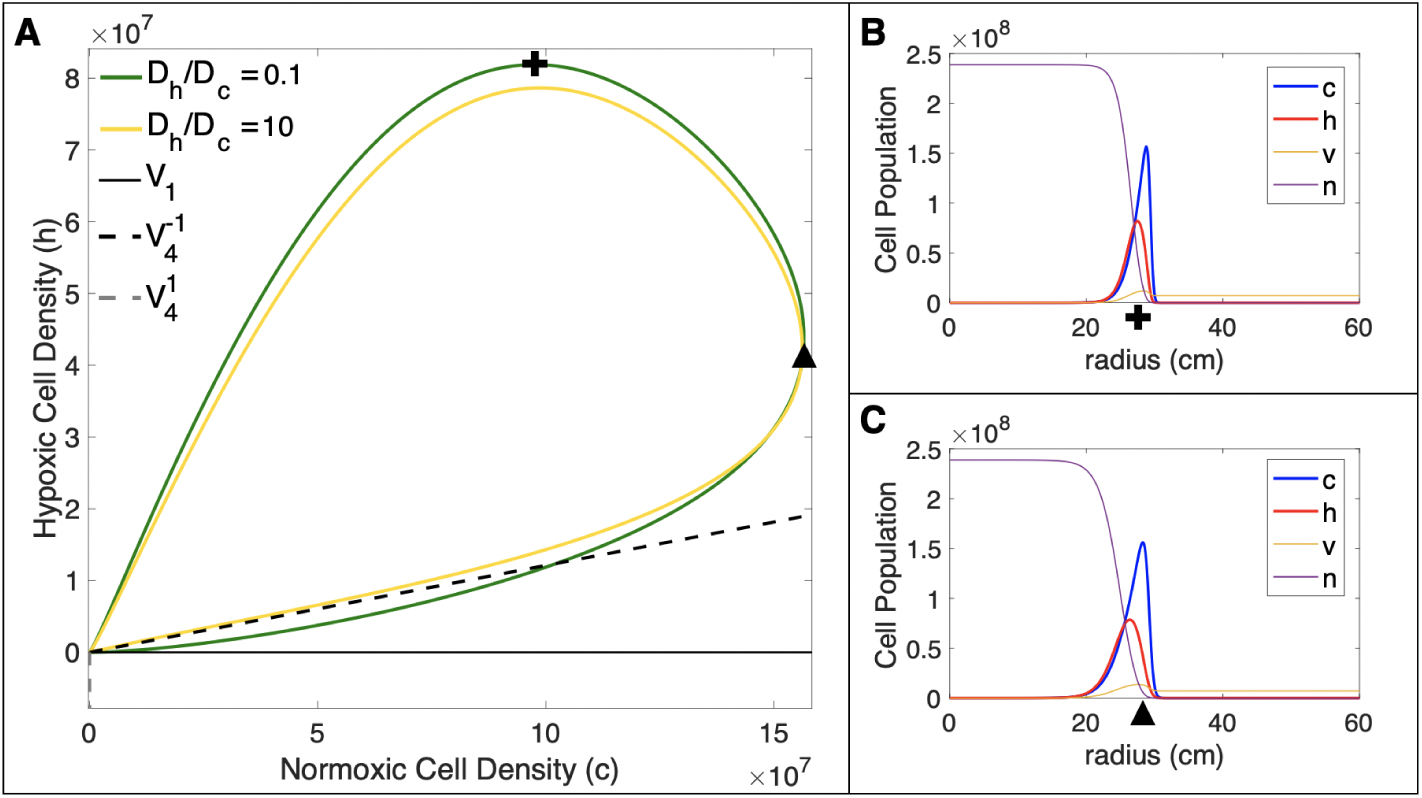
(A) We show the normoxic and hypoxic cell densities across space for a fixed snap-shot in time. For *D*_*h*_*/D*_*c*_ = 10^−1^, we see the dynamics follow *V*_1_ near (*c, h*) = (0, 0), which corresponds to the predicted minimum wave-speed, *s*_*min*_. For *D*_*h*_*/D*_*c*_ = 10^1^, the dynamics have shifted towards *V*_4_, corresponding with the faster numerical wave-speed we have observed. Here *Dc* = 10^1.5^mm^2^/year, *ρ* = 10^1.5^/year. Eigenvectors and simulations are shown for large simulated T2 sizes, to ensure convergence of numerical eigenvectors (29.5 - 30cm T2 radius, with corresponding time points of 1421 and 1910 days for *V*_4_). (B) The snap-shot of the simulation with *D*_*h*_*/D*_*c*_ = 0.1 used for subfigure A. The cross corresponds with the cross on subfigure A. (C) Corresponding snapshot with *D*_*h*_*/D*_*c*_ = 10, with the triangle corresponding with the triangle on subfigure A.

To provide further evidence concerning the traveling wave trajectory, we compared the gradient of the log of normoxic cells (*c*) with the eigenvalues *λ*_1_ and *λ*_4_. We see that for low values of *D*_*h*_*/D*_*c*_, the gradient more closely follows *λ*_1_ and for large values of *D*_*h*_*/D*_*c*_, the gradient closely follows *λ*_4_. We present examples of these results in the Appendix (Figure 6).

**Fig. 4.**
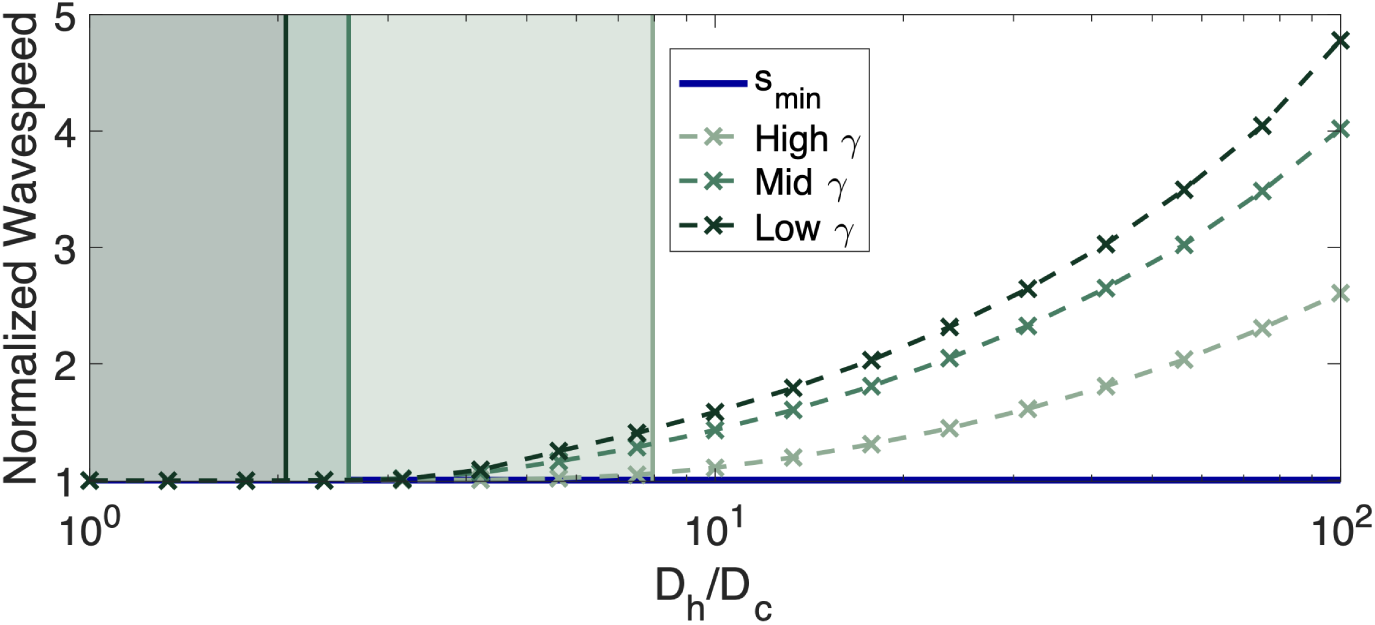
As *D*_*h*_*/D*_*c*_ is increased for varying *γ* values, we see an increase in the converged numerical wave-speed that is more pronounced for smaller values of *γ*; the corresponding thresholds *k* for wave-speeds faster than *s*_*min*_ are indicated. Wave-speeds taken as the average speed between 8 and 8.5cm of growth on simulated T2 MRI (16% total cell density threshold) and presented relative to *s*_*min*_.

**Fig. 5.**
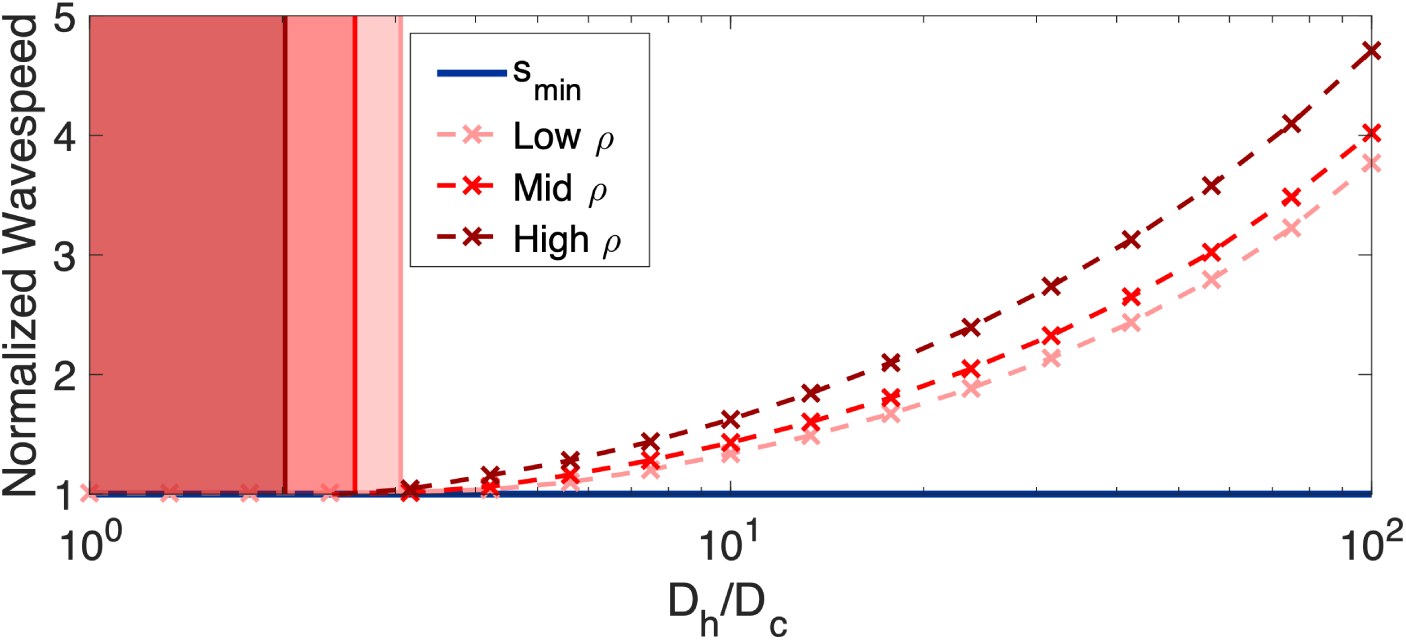
As *D*_*h*_*/D*_*c*_ is increased for varying *D*_*c*_ and *ρ* values, we see an increase in the relative wave-speed *s/s*_*min*_ that is more pronounced for larger values of *ρ*. The observed trend agrees with the expectation given by the corresponding values of *k*, which are also indicated. Wave-speeds taken as the average speed between 8 and 8.5cm of growth on simulated T2 MRI (16% total cell density threshold) and normalized against *s*_*min*_.

**Fig. 6.**
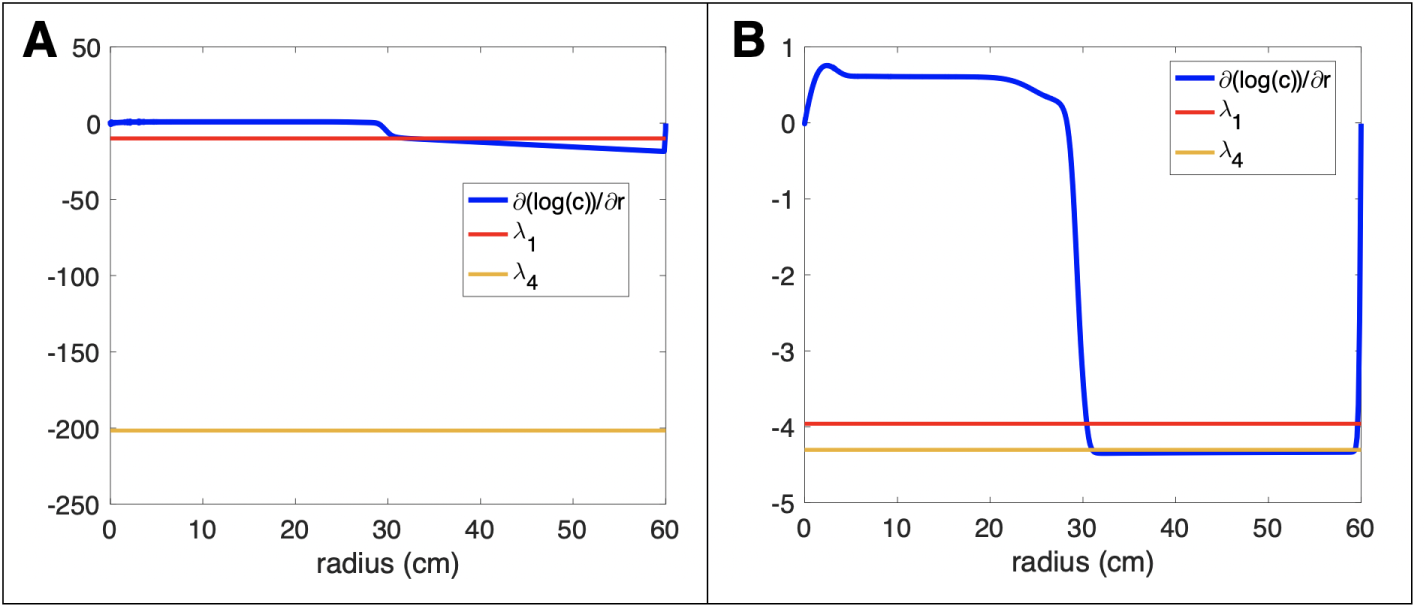
The gradient of the log of the normoxic cells is plotted for a T2 radius of 30cm. As described in the main text, the leading edge of this simulated gradient (ignoring boundary effects present close to the edge of the domain) should follow the eigenvalue that controls the dynamics of the PIHNA model. Simulations presented here correspond with those presented in Figure 3. The simulation with *D*_*h*_*/D*_*c*_ = 0.1 agrees more closely with *λ*_1_, whereas the simulation with *D*_*h*_*/D*_*c*_ = 10 follows *λ*_4_. The results of these support the eigenvalue and eigenvector analysis in the main body of this work.

### 5.2 Low switching rate from hypoxia to normoxia, *γ*, amplifies wave-speed increase for large values of *D*_*h*_*/D*_*c*_

The switching rate from a normoxic cell to a hypoxic cell (*β*) is not present in the eigenvalues that dominate the behavior of the wave-speed, nor in the expression for *k*. We do however note that the switching rate from a hypoxic cell phenotype back to a normoxic cell phenotype, *γ*, is present in the expression for *λ*_4_ (Equation (22)) and subsequently in the expression for the *D*_*h*_*/D*_*c*_ threshold, *k* (Equation (26)). We ran a similar set of simulations as in Section 5.1 with a higher value of *γ* = 0.5 and a lower value of *γ* = 0.005 to verify that the wave-speed increase, relative to *s*_*min*_, would be affected for varying *γ*. As expected, higher values of *γ* increase *k* and correspond to a lower wave-speed for equivalent *D*_*h*_*/D*_*c*_ values. We present these wave-speed results in Figure 4 where we also mark the corresponding values of *k*. For *γ* = 0.005, 0.05, and 0.5/day, we find *k* = 2.06, 2.60 and 7.95, respectively.

### 5.3 Wave-speed increase is more pronounced for faster-proliferating tumors

We also varied *ρ* to explore its effects on the increase in wave-speed for large values of *D*_*h*_*/D*_*c*_. We chose two more values of *ρ* = 10^1.25^/year (lower *ρ*), and *ρ* = 10^2^/year (higher *ρ*) and refer to the previous simulations with *ρ* = 10^1.5^/year as a mid-range *ρ*. Throughout all simulations, we set *D*_*c*_ = 10^1.5^mm^2^/year and *γ* = 0.05/day, leading Equation (26) to give threshold values of *k* = 2.19, 2.60 and 3.06 for higher, medium and lower *ρ* simulations, respectively.

We present the wave-speeds normalized against their predicted values of *s*_*min*_ (Equation (21)) in order to compare the simulation results across different values of *ρ*. We see for values of *D*_*h*_*/D*_*c*_ below their respective thresholds that the wave-speeds all follow their predicted minimum values. For simulations where *D*_*h*_*/D*_*c*_ is above the respective threshold *k*, we see an increased wave-speed, as expected (see Figure 5). This relative increase in wave-speed is more pronounced for larger values of *ρ*. Simulated tumors with larger *ρ* values are also faster-growing tumors as they have a faster minimum wave-speed (Equation (21)).

## Discussion

We have found an expression for the minimum wave-speed for the PIHNA model given by

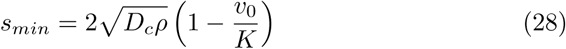

and shown that this predicted wave-speed is attained when normoxic cell diffusion is greater than or equal to the diffusion of hypoxic cells. We therefore have shown that the predicted minimum wave-speed is valid for previous publications of the PIHNA model [20]. However, due to the *in vitro* evidence indicating that hypoxia can increase migration [7, 23], we are interested in increasing the migration rate of hypoxic cells compared with normoxic cells in our extension of the PIHNA model. In the case that the hypoxic cells diffuse sufficiently faster than the normoxic cells, we see that the outward growth of the tumor is faster than the predicted minimum wave-speed value. In fact, we have quantified the value at which these faster rates of growth can occur through the threshold

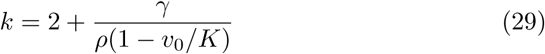

and note that the hypoxic cell diffusion has to be at least twice as fast as the normoxic cell diffusion. The threshold of hypoxic to normoxic cell migration rates is increased if the hypoxic cells can easily convert back to normoxia, and decreased for faster proliferating normoxic cell populations. This result suggests that faster-proliferating tumors that can only slowly recover from hypoxia are pushed to grow even faster by a highly migratory hypoxic sub-population, more so than slower-proliferating tumors that can easily recover from hypoxia.

The analysis presented here shows that the wave-speed dynamics do not depend on the vascular efficiency term, *V*, as long as *V* = 1 ahead of the wave. We also do not see a dependence on the switching rate from the normoxic cell density to the hypoxic cell density, *β*.

Mathematically, the increase in simulated wave-speed corresponds to a change in the asymptotic traveling wave trajectory as *D*_*h*_*/D*_*c*_ is increased, which causes an eigenvector associated with the hypoxic cell density characteristics to dominate the behavior of the PIHNA model. Biologically, this suggests that the faster migration of hypoxic cells can drive the growth of the whole tumor, as they migrate towards nutrient-rich environments and convert back to normoxic cells. If this conversion rate is high, the model suggests that the outward growth rate of the whole tumor is lower. The model does not predict that the wave-speed is affected by the proportions of hypoxic and normoxic cells. However, a reduction in vasculature ahead of the wave (*v*_0_) does increase the invasion speed of the tumor due to the appearance of *v*_0_ in the minimum predicted wave-speed. It would be interesting in future work to see how including a normal cell density affects these dynamics.

## A Appendix

As discussed in the main body of this work, we wanted to show that our PIHNA simulations were following different eigenvalues depending on the value of *D*_*h*_*/D*_*c*_. In the linearized regime ahead of the wave, we expect that 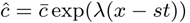, such that

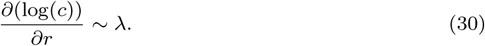

In Figure 6 we present simulations at a T2 radius of 30cm. We see for the simulation with *D*_*h*_*/D*_*c*_ = 0.1, *∂*(log(*c*))*/∂r* follows *λ*_1_ ahead of the traveling wave, whereas for *D*_*h*_*/D*_*c*_ = 10, *∂*(log(*c*))*/∂r* follows *λ*_4_.

## References

1. D.J. Brat, A.A. Castellano-Sanchez, S.B. Hunter, M. Pecot, C. Cohen, E.H. Hammond, S.N Devi, B. Kaur, and E.G. Van Meir. Pseudopalisades in glioblastoma are hypoxic, express extracellular matrix proteases, and are formed by an actively migrating cell population. Cancer Research, 64(3):920–927, 2004.

2. R.A. Fisher. The wave of advance of advantageous genes. Annals of Eugenics, 7(4):355–369, 1937.

3. A. Giese, R. Bjerkvig, M.E. Berens, and M. Westphal. Cost of migration: invasion of malignant gliomas and implications for treatment. Journal of Clinical Oncology, 21(8):1624–1636, 2003.

4. J.D. Gordan and M.C. Simon. Hypoxia-inducible factors: central regulators of the tumor phenotype. Current Opinion in Genetics & Development, 17(1):71–77, 2007.

5. H.L.P. Harpold, E.C. Alvord, and K.R. Swanson. The evolution of mathematical modeling of glioma proliferation and invasion. Journal of Neuropathology & Experimental Neurology, 66(1):1–9, 2007.

6. A. Hawkins-Daarud, R.C. Rockne, A.R.A. Anderson, and K.R. Swanson. Modeling tumor-associated edema in gliomas during anti-angiogenic therapy and its impact on imageable tumor. Frontiers in Oncology, 3:66, 2013.

7. M. Keunen, O.and Johansson, A. Oudin, M. Sanzey, S.A.A. Rahim, F. Fack, F. Thorsen, T. Taxt, M. Bartos, R. Jirik, et al. Anti-vegf treatment reduces blood supply and increases tumor cell invasion in glioblastoma. Proceedings of the National Academy of Sciences, 108(9):3749–3754, 2011.

8. P. Korkolopoulou, E. Patsouris, A.E. Konstantinidou, P.M. Pavlopoulos, N. Kavantzas, E. Boviatsis, I. Thymara, M. Perdiki, E. Thomas-Tsagli, D. Angelidakis, et al. Hypoxia-inducible factor 1*α*/vascular endothelial growth factor axis in astrocytomas. associations with microvessel morphometry, proliferation and prognosis. Neuropathology and Applied Neurobiology, 30(3):267–278, 2004.

9. D. Louis, H. Ohgaki, O. Wiestler, and W. Cavenee. WHO Classification of Tumours of the Central Nervous System, Revised. Fourth Edition. International Agency for Research on Cancer, 2016.

10. A. Martínez-González, G.F. Calvo, L.A.P. Romasanta, and V.M. Pérez-García. Hypoxic cell waves around necrotic cores in glioblastoma: a biomathematical model and its therapeutic implications. Bulletin of Mathematical Biology, 74(12):2875–2896, 2012.

11. S.C. Massey, M.C. Assanah, K.A. Lopez, P. Canoll, and K.R. Swanson. Glial progenitor cell recruitment drives aggressive glioma growth: mathematical and experimental modelling. Journal of The Royal Society Interface, page rsif20120030, 2012.

12. A. Roniotis, V. Sakkalis, E. Tzamali, G. Tzedakis, M. Zervakis, and K. Marias. Solving the pihna model while accounting for radiotherapy. In Advanced Research Workshop on In Silico Oncology and Cancer Investigation-The TUMOR Project Workshop (IAR-WISOCI), 2012 5th International, pages 1–4. IEEE, 2012.

13. D.L. Silbergeld and M.R. Chicoine. Isolation and characterization of human malignant glioma cells from histologically normal brain. Journal of Neurosurgery, 86(3):525–531, 1997.

14. R. Stupp, M.E. Hegi, W.P. Mason, M.J. van den Bent, M.J.B. Taphoorn, R.C. Janzer, S.K. Ludwin, A. Allgeier, B. Fisher, K. Belanger, et al. Effects of radiotherapy with concomitant and adjuvant temozolomide versus radiotherapy alone on survival in glioblastoma in a randomised phase iii study: 5-year analysis of the eortc-ncic trial. The Lancet Oncology, 10(5):459–466, 2009.

15. R. Stupp, W.P. Mason, M.J. Van Den Bent, M. Weller, B. Fisher, M.J.B. Taphoorn, K. Belanger, A.A. Brandes, C. Marosi, U. Bogdahn, et al. Radiotherapy plus concomitant and adjuvant temozolomide for glioblastoma. New England Journal of Medicine, 352(10):987–996, 2005.

16. A. Swan, T. Hillen, J.C. Bowman, and A.D. Murtha. A patient-specific anisotropic diffusion model for brain tumour spread. Bulletin of Mathematical Biology, 80(5):1259–1291, 2018.

17. K.R. Swanson. Mathematical modeling of the growth and control of tumors, 1999.

18. K.R. Swanson, E.C. Alvord, Jr, and J.D. Murray. A quantitative model for differential motility of gliomas in grey and white matter. Cell Prolif, 33(5):317–29, Oct 2000.

19. K.R. Swanson, C. Bridge, J.D. Murray, and E.C. Alvord. Virtual and real brain tumors: using mathematical modeling to quantify glioma growth and invasion. Journal of the Neurological Sciences, 216(1):1–10, 2003.

20. K.R. Swanson, R.C. Rockne, J. Claridge, M.A.J. Chaplain, E.C. Alvord, and A.R.A. Anderson. Quantifying the role of angiogenesis in malignant progression of gliomas: in silico modeling integrates imaging and histology. Cancer Research, 71(24):7366–7375, 2011.

21. K.R. Swanson, R.C. Rostomily, and E.C. Alvord. A mathematical modelling tool for predicting survival of individual patients following resection of glioblastoma: a proof of principle. British Journal of Cancer, 98(1):113–119, 2008.

22. Christina H Wang, Jason K Rockhill, Maciej Mrugala, Danielle L Peacock, Albert Lai, Katy Jusenius, Joanna M Wardlaw, Timothy Cloughesy, Alexander M Spence, Russ Rockne, et al. Prognostic significance of growth kinetics in newly diagnosed glioblastomas revealed by combining serial imaging with a novel biomathematical model. Cancer research, 69(23):9133–9140, 2009.

23. David Zagzag, Yevgeniy Lukyanov, Li Lan, M Aktar Ali, Mine Esencay, Olga Mendez, Herman Yee, Evelyn B Voura, and Elizabeth W Newcomb. Hypoxia-inducible factor 1 and vegf upregulate cxcr4 in glioblastoma: implications for angiogenesis and glioma cell invasion. Laboratory Investigation, 86(12):1221, 2006.

24. David Zagzag, Hua Zhong, Joanne M Scalzitti, Erik Laughner, Jonathan W Simons, and Gregg L Semenza. Expression of hypoxia-inducible factor 1*α* in brain tumors. Cancer, 88(11):2606–2618, 2000.

